# Structural basis for selective remdesivir incorporation by SARS-CoV-2 RNA polymerase, and S759A resistance

**DOI:** 10.1101/2025.09.04.674295

**Authors:** Calvin J. Gordon, Mizar F. Oliva, Hery W. Lee, Quinten Goovaerts, Brent De Wijngaert, Matthias Götte, Kalyan Das

## Abstract

Nucleoside analogs (NAs) have been successfully used to treat viral infections. dNTP analogs are primarily DNA chain terminators, while NTP analogs, such as remdesivir, can inhibit as delayed chain terminators or when in the template strand. Determining the frequency of remdesivir triphosphate (RTP) incorporation in the presence of the competing ATP can help in understanding different modes of viral RNA-dependent RNA polymerase (RdRp) inhibition by NTP analogs. We employed enzymatic assays, mass spectrometry, and cryo-EM to show that SARS-CoV-2 RdRp preferentially incorporates RTP, outcompeting 10-fold higher ATP concentration; however, successive RTP incorporations are less favoured when ATP is present. Structures of SARS-CoV-2 RdRp in this and previous studies demonstrate resilience of remdesivir:UMP base pair to translocation, explaining the reduced preference for conjugate incorporations. Together, the RTP versus ATP incorporation is driven by their relative concentration and structural rigidity of remdesivir:UMP, ultimately limiting the number of incorporated remdesivir in a fully synthesized RNA strand. The S759A mutant confers RTP resistance, and the structures of S759A RdRp catalytic complexes reveal that altered ribose-ring conformation and repositioning of the primer 3′-end nucleotide contribute to RTP resistance. These findings enhance our understanding of non-obligate NTP analogs and provide insight into S759A resistance mechanism.

## Introduction

Nucleoside/nucleotide analogs (NAs) constitute an important class of antiviral drugs used in the treatment of HIV, hepatitis B virus (HBV), cytomegalovirus (CMV), herpes simplex virus (HSV), hepatitis C virus (HCV), and severe acute respiratory syndrome coronavirus 2 (SARS-CoV-2) infections. In their triphosphate (TP) form, these drugs act as substrates for viral polymerases, which catalytically incorporate NAs into the primer strand, resulting in inhibition of replication and/or transcription of the viral genome. However, the potency, specificity, and mechanisms of action can differ among NAs due to their chemical modifications relative to the natural NTP/dNTP substrates. dNTP analogs such as zidovudine (AZT), lamivudine (3TC), and tenofovir (TFV) target viral DNA polymerase or reverse transcriptase (RT), acting as obligate DNA chain terminators^1,2^; *i.e.* these inhibitors lack the 3′-OH group and prevent incorporation of the next nucleotide. However, the development of NAs against RNA viruses has been more challenging, and the approved NTP analog prodrugs, such as sofosbuvir (SOF), remdesivir (RDV), and molnupiravir (MOV), feature a 3′-OH group. While SOF has been developed for the treatment of HCV infection, RDV and MOV show a broader spectrum of antiviral activity and have been approved in several jurisdictions for the treatment of COVID-19. Only sofosbuvir prevents the incorporation of the next nucleotide by HCV RdRp; therefore, functioning as a non-obligate RNA immediate chain terminator^3^. Conversely, RDV and MOV do not block RNA synthesis immediately following incorporation^4,5^. This is important for RNA viruses with proofreading activity. Coronaviruses, such as SARS-CoV-2, possess a 3′-to-5′ proof-reading exoribonuclease able to remove mismatched nucleotides or chain-terminating nucleotide analogs^6–9^. RDV and molnupiravir exhibit different mechanisms of inhibition that help to evade proofreading^10–12^.

Molnupiravir is incorporated as a CTP or UTP analog and does not show significant inhibition of RNA synthesis. Instead, its antiviral effect is mediated when embedded in the template strand, where it permits the incorporation of ATP and GTP, resulting in erroneous viral RNAs^13–15^. RDV is a *C*-adenosine 1′-cyano-modified monophosphate prodrug that demonstrated broad antiviral efficacy against various positive-sense RNA viruses, such as coronaviruses and enteroviruses^16,17^, as well as nonsegmented negative-sense RNA viruses like pneumoviruses and paramyxoviruses^18^. However, reduced potency has been observed against segmented RNA viruses like orthomyxoviruses^18,19^. Understanding the mechanism of RDV incorporation over the natural nucleoside AMP, the structural basis of RDV inhibition, and the molecular mechanism of drug resistance would help design improved NAs targeting viral RNA-dependent RNA polymerases (RdRps), including that of SARS-CoV-2.

The heterotrimeric SARS-CoV-2 RdRp complex, composed of the non-structural proteins Nsp7, Nsp8, and Nsp12, is responsible for viral RNA synthesis. Template binding and nucleotide incorporation are predominantly driven by the catalytic motifs, often referred to as A through G, which are contained within Nsp12. These motifs are sequence-derived and well-conserved among RdRps from diverse RNA viruses^20^. Nsp7 and Nsp8 appear to play structural roles. Kinetic and biochemical studies have demonstrated that remdesivir triphosphate (RTP) is a better substrate than ATP, its natural counterpart^21,22^. Cryogenic-electron microscopy (Cryo-EM) structures of the SARS-CoV-2 RdRp active complex with RTP incorporated^23^ or bound as a substrate^24^ have revealed the molecular interactions and provided the structural basis for compound recognition. RDV functions as a delayed chain terminator, and the current understanding is that inhibition occurs three nucleotides after the first RTP is incorporated^22,25,26^. Several biochemical, structural, and mutagenesis studies support a mechanism that disfavors the translocation of the dsRNA containing remdesivir monophosphate (RMP), identifying a steric clash between the 1′-cyano group of RDV in the primer strand and Nsp12 residue S861 as the limiting factor^22,25–28^. However, this translocation barrier can be easily overcome with increased NTP concentrations that better mimic cellular conditions^10,22^. Thus, it is plausible that RMPs become embedded in the synthesized viral genomes and could serve as a templating base for subsequent rounds of replication. Indeed, the incorporation of UTP opposite RMP is compromised, resulting in template-dependent inhibition^27,29^. It remains unclear when and how RDV outcompetes ATP during RNA strand synthesis, and whether it is possible to predict the frequency and specific sites of RMP in the template strand.

We employed biochemical and mass spectrometry studies and determined cryo-EM structures of the SARS-CoV-2 RdRp/dsRNA catalytic complexes with ATP and RTP, to understand the preference of RTP versus ATP incorporation during RNA synthesis. We show that RTP is preferentially incorporated at the first site even in the presence of ATP at 10-fold excess. Additionally, we show that consecutive incorporations are influenced by the relative concentration of the analog versus ATP. The cryo-EM structures of SARS-CoV-2 RdRp/dsRNA following consecutive RTP or ATP incorporations and demonstrate that the structural rigidity of the RMP:UMP (R:U) base pair (bp) impairs the translocation efficiency of dsRNA through the RNA-binding cleft of the RdRp. Additionally, biochemical studies and structures of Nsp12- S759A mutant RdRp complexes with RTP or ATP were conducted to elucidate the mechanism by which the mutation confers drug resistance. The study on the Nsp12-S759A mutant indicates that the altered ribose-ring conformation, repositioning of the primer 3′-end at the P-site, and loss of the side-chain interactions influence substrate binding, incorporation, and translocation characteristics of the mutant enzyme.

## Results

### Competition between ATP and RTP

Earlier studies have examined the preferential incorporation of RTP over ATP by the SARS-CoV-2 RdRp, either with one or the other, but not when both are present and competing in the reaction mixture. Individually, RTP is incorporated 2 to 3 times more efficiently than ATP^21,22^. One such study was conducted under pre-steady-state conditions using a dsRNA composed of a 40-mer template and a 6-carboxyfluorescein (FAM)-labelled 20-mer primer that represents the 3′-end of the SARS-CoV-2 genome preceding the poly(A) tail^21^. Importantly, this sequence contains a homopolymeric stretch of uridines that can facilitate the sequential incorporation of four RTP or ATP molecules (Figure 1A, *top*). The study found that, while the first RTP molecule is incorporated more efficiently, the second and third sequential RTP incorporations are less efficient. However, treating a SARS-CoV-2-infected cell means that viral RNA synthesis occurs in the presence of both ATP and RTP, and RTP competes with excess ATP for binding and incorporation. Therefore, we hypothesized that the relative concentration of RTP to ATP is likely to influence which one is incorporated, and there may be a site preference for one over the other.

**Figure 1:**
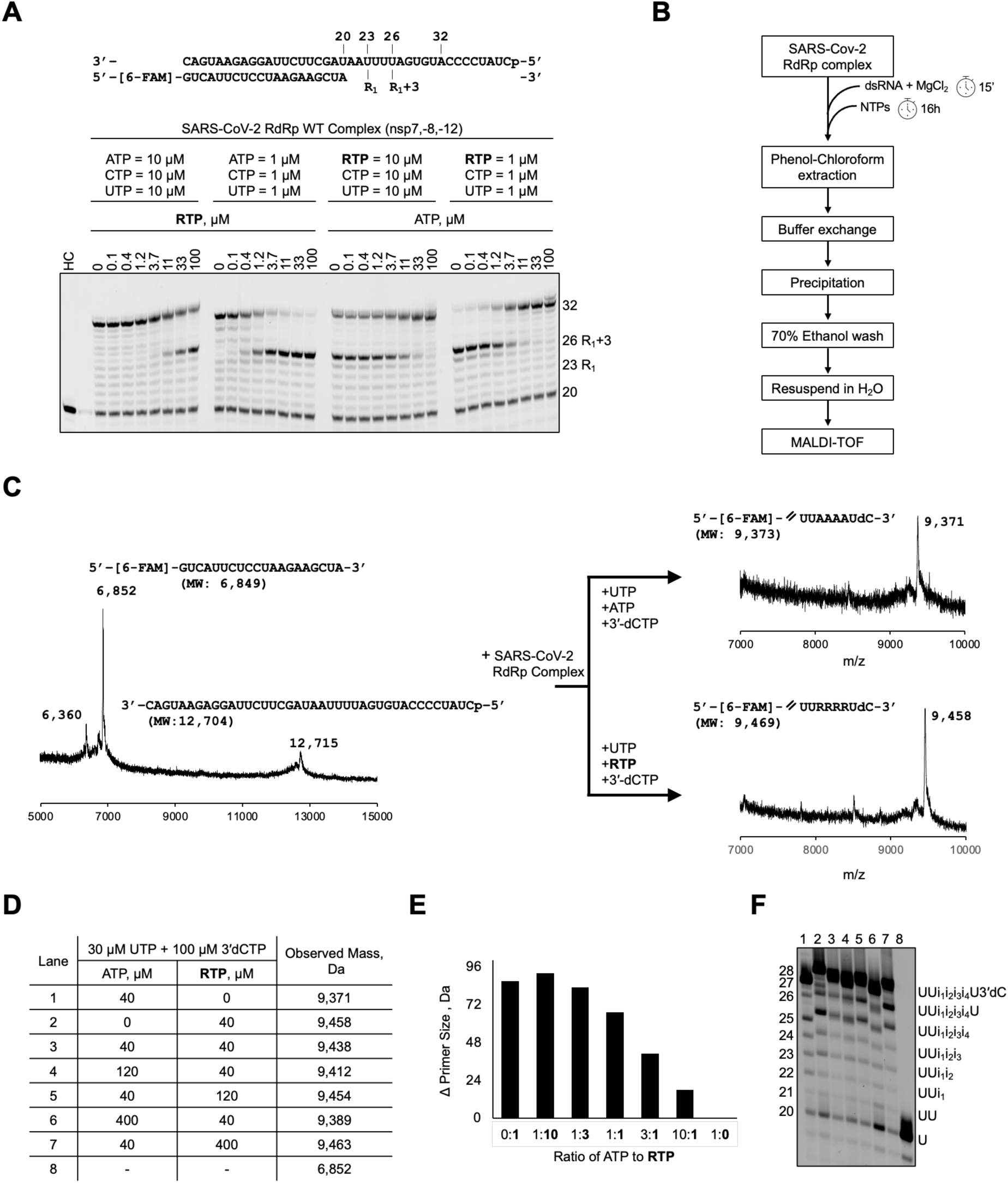
Competition between ATP and RTP for incorporation. **A**, RNA synthesis was evaluated on a dsRNA oligo that consists of a 40-nt template strand annealed to a 20-nt primer strand (*top*). Here, a stretch of uridines spans the templated positions 23 through 26. The pattern of SARS-CoV-2-catalyzed RNA synthesis in the presence of increasing concentrations of RTP (left) or ATP (right). When ATP concentration is reduced or RTP is increased, product formation at position 26 (i-3) becomes more prominent, consistent with delayed chain termination when RTP is incorporated at position 23 (i) and translocated to i-3. **B**, Schematic depicting the method used to prepare SARS-CoV-2-catalyzed RNA primers for analysis by MALDI-TOF mass spectrometry (MS). **C**, MALDI-TOF mass spectrum of the 20- nt RNA primer-strand and the 40-nt template-strand (*left*); their expected mass is shown in parentheses below the sequence. Polymerase extension reactions were performed by incubating UTP, 3’-dCTP, and ATP (*right, top*) or RTP (*right, bottom*) with the SARS-CoV-2 RdRp complex. The sequence of the extended portion of the RNA primer is shown above the graph; the size, indicated below in parenthesis, corresponds to the entire 28-nt RNA product. **D**, the nucleotide concentrations that were supplemented to the SARS-CoV-2 RdRp complex for RNA synthesis (*columns 2 and 3*), and the mass of the RNA was determining by mass spectrometry (*column 4*). The purified RNA product was analysed using MALDI-TOF MS with a 2,4,6-trihydroxyacetophenone (THAP) matrix. **E**, the difference in observed masses of different RNA primers synthesized in the presence of varying concentrations of RTP and ATP. The resulting RNA products generated from the conditions are listed in **D**. **F**, RNA products corresponding to conditions 1 – 8 in **D** are shown in 20% UREA-PAGE. On the right side of the gel are the likely products generated; “i” represents either an ATP or RTP incorporation event. RTP is marginally larger than ATP and migrates slowly. Therefore, a shift in size at positions 23 to 26 between the different reaction conditions is noticeable for RTP-incorporated RNAs (in lanes 1 vs 2 and 6 vs 7).

In this study, we used the same 40/20-nt template/primer dsRNA that can support the incorporation of four conjugate RTP molecules, four conjugate ATP molecules, or a combination of both. We evaluated the pattern of RNA synthesis in response to different ATP to RTP concentrations (Figure 1A, *bottom*). Additionally, 10 µM of UTP and CTP were supplemented to the reaction to support the synthesis of a 32-nt RNA product. We observed that the first RTP incorporation at position 23 results in an intermediate product at position 26, consistent with the delayed chain termination mechanism described earlier. However, additional pause site(s), corresponding to subsequent RTP incorporation, could not be identified. This suggests that, although the template sequence can support multiple RTP incorporations, multiple consecutive RTP incorporations may not occur even at higher RTP concentrations compared to ATP and/or the delayed chain-termination event at the corresponding third upstream position does not occur for subsequently incorporated RTPs. The 24 Da mass difference between RTP and ATP, Δ(RTP-ATP), is difficult to resolve using a traditional gel-based assay. Therefore, to determine whether RTP or ATP is incorporated at positions 24 to 26 following the first RTP incorporation at position 23, we evaluated the size of the RNA primer strand using a systematic mass spectrometry study as outlined below. In this experiment, the SARS-CoV-2 RdRp carried out RNA synthesis in the presence of ATP and RTP at different molar ratios.

Building on the above gel-based assay, we first optimized RNA synthesis conditions to achieve the desired 28-mer dsRNA at a high yield required for mass spectrometry analysis (Figure 1B, see Methods). Here, CTP was substituted with 3′-dCTP to avoid potential misincorporation and to ensure the runoff RNA products were not longer than 28 nucleotides, thereby simplifying the readout to focus on the homopolymeric region of interest. The RNA was isolated via phenol-chloroform extraction, followed by precipitation in sodium acetate and isopropanol. The purified RNA product was then subjected to Matrix-Assisted Laser Desorption/Ionization Time-of-Flight (MALDI-TOF) mass spectrometry to determine its mass. Processing the labelled dsRNA substrate alone resulted in a spectrum with a single primary peak at a m/z of 6,852 Da, 3 Da smaller than the predicted mass of the labelled 20-mer primer (Figure 1C, *left*). Conversely, two less intense peaks are observed at m/z values of 6,360 Da and 12,715 Da. The 6,360 Da peak presumably represents a population of unlabeled RNA primer or a degraded product. The 12,715 Da peak corresponds to the RNA template, which has a predicted mass of 12,704 Da. Extension of the 20-mer RNA primer with four ATP or RTP produced oligos with observed masses of 9,371 Da or 9,458 Da, respectively, which are close to their expected masses for a 28-nt product. (Figure 1C, *right*). A mass difference of 87 Da between the two RNA products is within 7 Da of the theoretical value of 4Δ(RTP-ATP). Subsequently, the 20-nt primers were extended to 28-nt RNAs under different competitive ATP to RTP ratios, and the masses of the primers were interpreted to evaluate the number of RTP incorporations (Figure 1D and E, Figure S1).

RTP was preferentially incorporated at the first site, even at an ATP to RTP ratio of 10:1; however, the three subsequent incorporations were ATP. Here, the observed RNA mass is higher by 1Δ(RTP-ATP) compared to the reference RNA with ATP. The faster migration of the RNA product at positions 24, 25, and 26, and a RTP-dependent pausing at 26 (Figure 1A) suggests that an RMP occupies only position 23 (Figure 1F, *lane 6*). At a 3:1 ATP to RTP ratio, the analog was preferentially incorporated at two positions, and when the ratio was 1:1, there were three RTP incorporations. The mass differences in all three cases are near multiples of ∼24 Da, Δ(RTP-ATP), suggesting that RTP incorporation is preferential rather than random. At ATP to RTP concentration of 1:3, only one or fewer AMP is incorporated, and at a 1:10 ratio of ATP to RTP, only RTP is incorporated at all four positions. It is important to note that our experiment does not show the preferred positions between sites 24 and 26 for RTP. Together, these data reveal that the incorporation of multiple RTP molecules is contingent upon the relative concentration of RTP to ATP. Considering that the cellular concentration of ATP is significantly higher than that of RTP, the inhibitor is preferentially incorporated at certain sites only. Additionally, the experimental platform presented in this study has broader implications for addressing the competitive incorporation of NAs versus their natural counterparts in other systems.

### Structures of wild-type and S759A SARS-CoV-2 RdRp complexes

To understand the molecular basis for the selective incorporation of RTP, we determined the structures of SARS-CoV-2 RdRp complexes incorporating RTP or ATP. In parallel, we also determined the analogous structure of the Nsp12-S759A mutant complex to understand the molecular mechanism of the resistance mutation (Figure 2). The same 40/20-nt dsRNA was used for the structural studies. The wild-type (WT) and mutant complexes were incubated with UTP and ATP to catalytically extend the primer by seven nucleotides, complementing the template overhang sequence “AAUUUUA”. The primer is extended by seven nucleotides, two UTP, four ATP, followed by one UTP, and the 3′-end of this elongated 27-nt primer is translocated to the P-site, leaving the empty N-site ready to bind a CTP that was not supplied (Figure 2B). In contrast, when incubated with UTP and RTP, both wild-type and the mutant complexes stalled after incorporation of five nucleotides, “UURRR”, and the 3′-end RMP was positioned at the pre-translocated N-site, preventing the binding of the next RTP (Figure 2C); identical ATP and RTP incorporations were also observed for the S759A mutant complex. The structures of the wild-type RdRp with ATP (WT-AMP) and RTP (WT-RMP), and the S759A mutant with ATP (S759A-AMP) and with RTP (S759A-RMP) were obtained at 2.8, 2.9, 3.1, and 2.9 Å resolution, respectively (Table S1, Figure S2).

**Figure 2:**
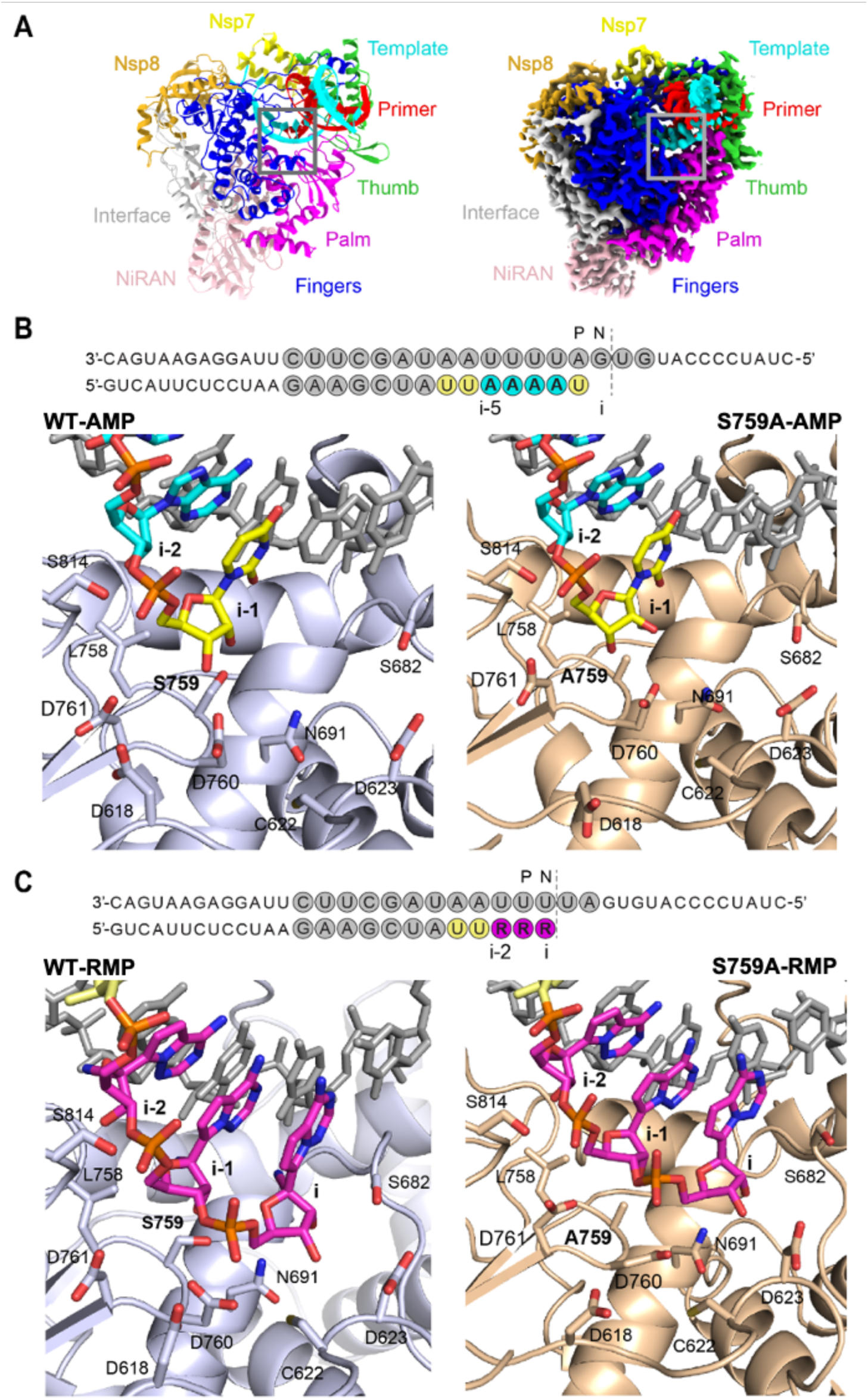
Structures of WT and S759A mutant RdRp complexes. **A**, Structural model (left) and cryo-EM density map (right) of the WT RdRp/RNA complex (WT-AMP). RdRp complex subunits and Nsp12 domains are colored as follows: Nsp7 (yellow); Nsp8 (gold); Nsp12 domains thumb (green), palm (magenta), fingers (blue), NiRAN (pink), interface (grey); RNA template (cyan) and primer (red) strands. Active site region is boxed in grey. **B**, Schematic representation of the dsRNA present in the WT-AMP and S759A-AMP structures. P and N sites are indicated above the sequence. The bases of template and primer strands are colored in grey and incorporated nucleotides in yellow (UMP) and cyan (AMP). The WT (light blue) and S759A (wheat) are shown in cartoon representation. **C**, Schematic representation of the dsRNA present in the WT-RMP and S759A-RMP structures, with protein color scheme and annotations as in **B**; RMP is in magenta. The key amino acid residues in the active site regions are shown as sticks in panels **B** and **C**.

Structures of the SARS-CoV-2 RdRp complex with incorporated RMP or incoming RTP have been reported^23–26,30^. For comparison of the structures, the upstream positions of the nucleotide analog are described in reference to the active site, denoted as position i. The published structures have RTP positioned at i, prior to incorporation (PDB: 7UO4) and after incorporation (PDB: 7BV2), incorporated RMP translocated by one nucleotide to i-1 position (PDB: 7C2K), and translocated by three nucleotides to i-3 position (PDB: 7B3B, 7B3C and 7L1F); the upstream n^th^ site is counted from i as i-n position (Figure 2B and C). The position i in the structure is also the NTP-binding site (N-site), the i-1 position is the post-translocated site (P- site). In comparison, our RMP complexes are stalled after three RTP incorporations, with RMPs occupying the i-2, i-1, and i positions of the primer; the experimental density for the RNA substrate unambiguously traced the sequence (Figures S3 and S4).

### Remdesivir introduces rigidity and impairs translocation

Our WT-RMP structure is in agreement with the previously described kinetic study, confirming the incorporation of three consecutive RMPs into the same RNA duplex^21^. Unlike a follow-up structural study on this dsRNA with four incorporated RMPs^26^, we did not observe the incorporation of the fourth RTP. Therefore, our structure, which captures RMP at the i-2 position, completes the sequence of structures with the first incorporated RMP occupying each position, starting from i-3 to i. Biochemically, the position of the first incorporated RMP is important for inhibition, which occurs at the i-3 position. In contrast, the structural data, including ours, suggest that RMP resists translocation from the i-th position itself; however, the presence of the next NTP helps drive the translocation^30^.

To assess the potential structural impacts of the RMP at different positions along the RNA-primer on the overall conformation of the RdRp complex, we aligned available structures based on respective Nsp12 C⍺ superpositions. The superposition of the four structures reported in this study align well with a root mean square deviation (RMSD) of ∼0.5 Å, indicating no significant overall difference. These structures also align well with earlier reported structures (PDB IDs: 7BV2, 7C2K, 7B3B, 7B3C, 7L1F); Nsp12 C⍺ superimpose with an RMSD of ∼0.5 Å for any pair of structures except for 7L1F, for which the RMSD is ∼1.2 Å. The low RMSD among the structures suggests a well-defined RNA-binding cleft of the RdRp that maintains its shape to guide the translocation of the RNA duplex rather than being perturbed in the translocation process.

It is reasonable to assume that the template-primer base-pairs (bps) require certain flexibility for translocating through the RNA-binding channel of the RdRp, and the presence of the 1′- cyano group alters the size and structural rigidity of the R:U bp compared to A:U^1^. For example, a nucleotide ribose ring in dsRNA favors a 3′-endo conformation; however, it can transiently switch to a 2′-endo conformation. In contrast, as observed in all structures, the ribose rings of RMPs in the RNA are presumably locked in a 3′-endo conformation, primarily restricted by the C1′-cyano group (Figure 3A). This locked 3′-endo conformation may also provide a lower entropic penalty for RTP binding at the N-site, a contributing factor for the higher selectivity of RTP over ATP. Following incorporation, the higher structural rigidity of R:U over A:U bp may be contributing to the lower translocation efficiency of R:U. To understand the implications of R:U bp at different positions of the dsRNA substrate, we evaluated the minor groove and backbone C1′-C1’ distances of each bp from positions i-5 to i between the WT-AMP and WT- RMP structures (Figure 3B). The RMP-incorporated structures that have actively translocated the dsRNA show an increase of the minor groove width around i-1 site (Figure 3C). For the backbone distances, the most significant difference is observed at position i-4, where the A:U bp exhibits a change in distance of about 0.5 Å at C1′-C1′ atoms compared to the RMP structures (Figure 3D). This may suggest that R:U bp at i-3 lacks the required flexibility for translocating to i-4, aligning with the delayed chain termination hypothesis; the structure with four incorporated RMPs to the same template/primer (PDB: 7I1F) appears to be an outlier from the remaining structures including ours. The minor groove widths in our RMP- and AMP- incorporated structures show two distinct trends, with little influence from the mutation (Figure 3E). However, the observed differences in the RNA tracks between the structures are not highly significant. Considering that the dsRNA-binding cleft is unaltered in the structures and dsRNA must wiggle for translocating through the cleft, a R:U base-pair with its restricted flexibility may experience hindrance in these dynamic steps of translocation.

**Figure 3:**
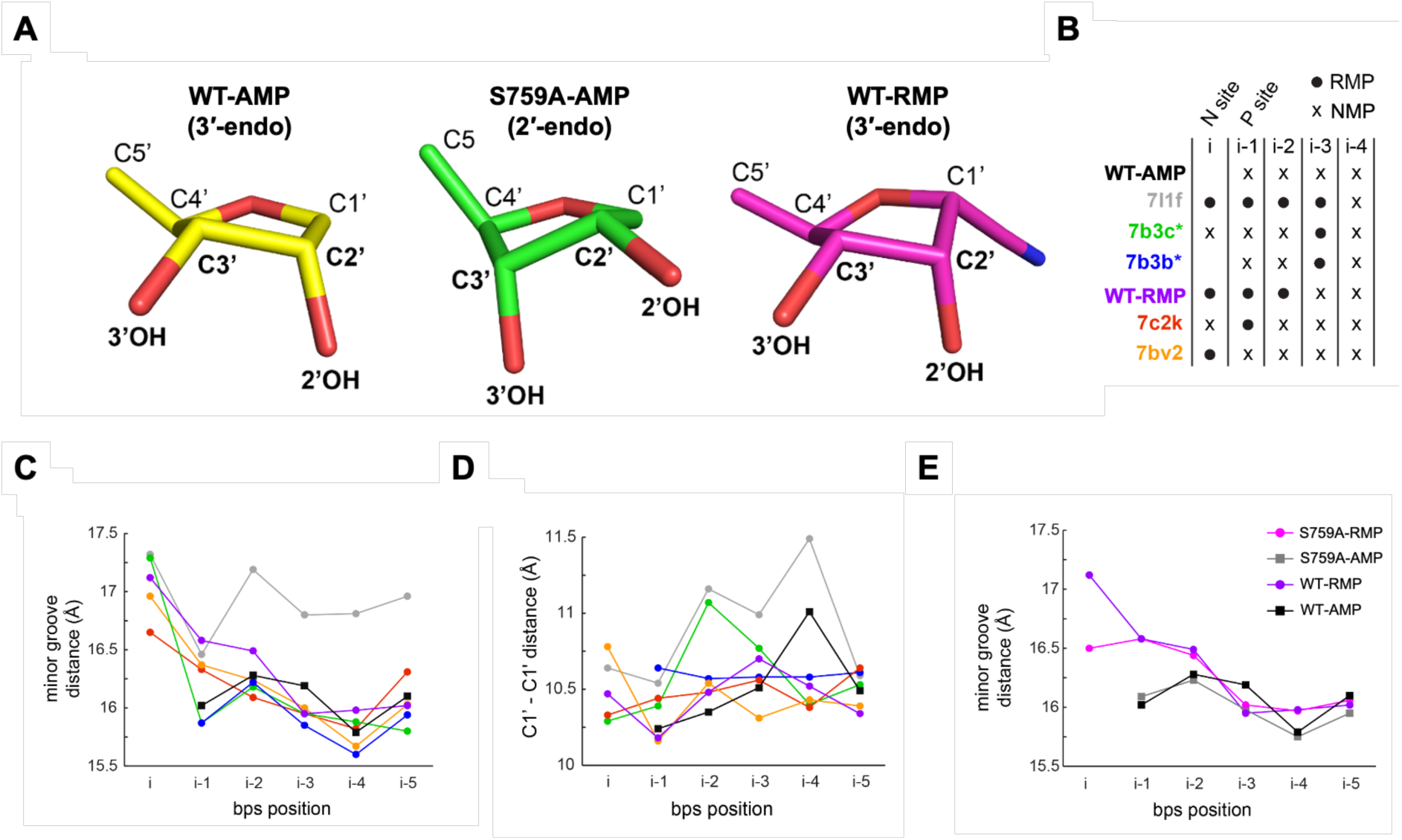
Effect of remdesivir incorporation on translocation and RNA rigidity. **A**, Conformations of the sugar rings of the last incorporated nucleotide occupying P-site for the WT-AMP (yellow) is 3′- endo and for S759A-AMP (green) is 2′-endo. The conformation of the sugar ring of all RMPs (magenta) structures is in 3-endo. **B**, The available SARS-CoV-2 RdRp structures containing remdesivir at different positions in the primer strand are indicated in the scheme by their PDB IDs and the structures reported in this study are labeled. Remdesivir (RMP) position is indicated as a black dot and AMP as an x. **C**, Minor groove distance is plotted for each bp position between i to i-5 for all the structures listed in the scheme in **B** with curves following the same color code. **D**, Distances between C1′-C1’ of bps are plotted for each position between I and i-5, curves follow the same color code of the scheme in **B**. **E**, Minor groove distances are plotted for each bp position between i to i-5 for all the structures obtained in this study.

### Successive incorporations of RMP are not favored

Biochemically, to ascertain that a pause occurs after three sequential RMP incorporation events, as observed in our structures, RNA synthesis by the SARS-CoV-2 RdRp complex was monitored on the labelled dsRNA duplex (Figure 4). Here, the RdRp complex was equilibrated with RNA substrate and MgCl_2_ before catalysis, which was initiated by the addition of NTPs. These reactions were performed in the presence of 4 mg/mL of heparin to ensure the RNA products were synthesized under single-turnover conditions^31^. The addition of 1 µM ATP and 1 µM UTP could readily generate the expected 27-mer RNA product, as CTP was not supplied in this experiment. Alternatively, substituting 1 µM RTP for ATP resulted in product formation at position 25 (i-2) within the first 30 seconds, after which a conversion to a 26-nt product occurred. This transition signifies the rapid incorporation of the first three RTP molecules, followed by a slowdown in RNA catalysis due to the inefficient incorporation of the fourth RTP molecule. Indeed, similar observations via capillary electrophoresis indicate that at 20 seconds, the fourth RTP was incorporated much more slowly than the first three^21,26^. Increasing RTP concentration to 10 µM accelerated the incorporation of the fourth RTP at position 26 within the first 30 seconds. Subsequently, the subtle formation of a 27-mer product was observed when the concentration of UTP was increased to 10 µM. Unlike the 25-mer product observed at 1 µM RTP and UTP, the larger 26-mer products could not be extended over time. The inefficient nucleotide incorporation following four RMPs agrees with the well-described mechanism of delayed-chain termination by RDV. The combination of our structural and biochemical data, along with previous kinetic data, suggests that the incorporation of the first three RMP is highly favoured. However, the incorporation of the next two nucleotides is possible when the RTP and/or NTP concentration is increased.

**Figure 4:**
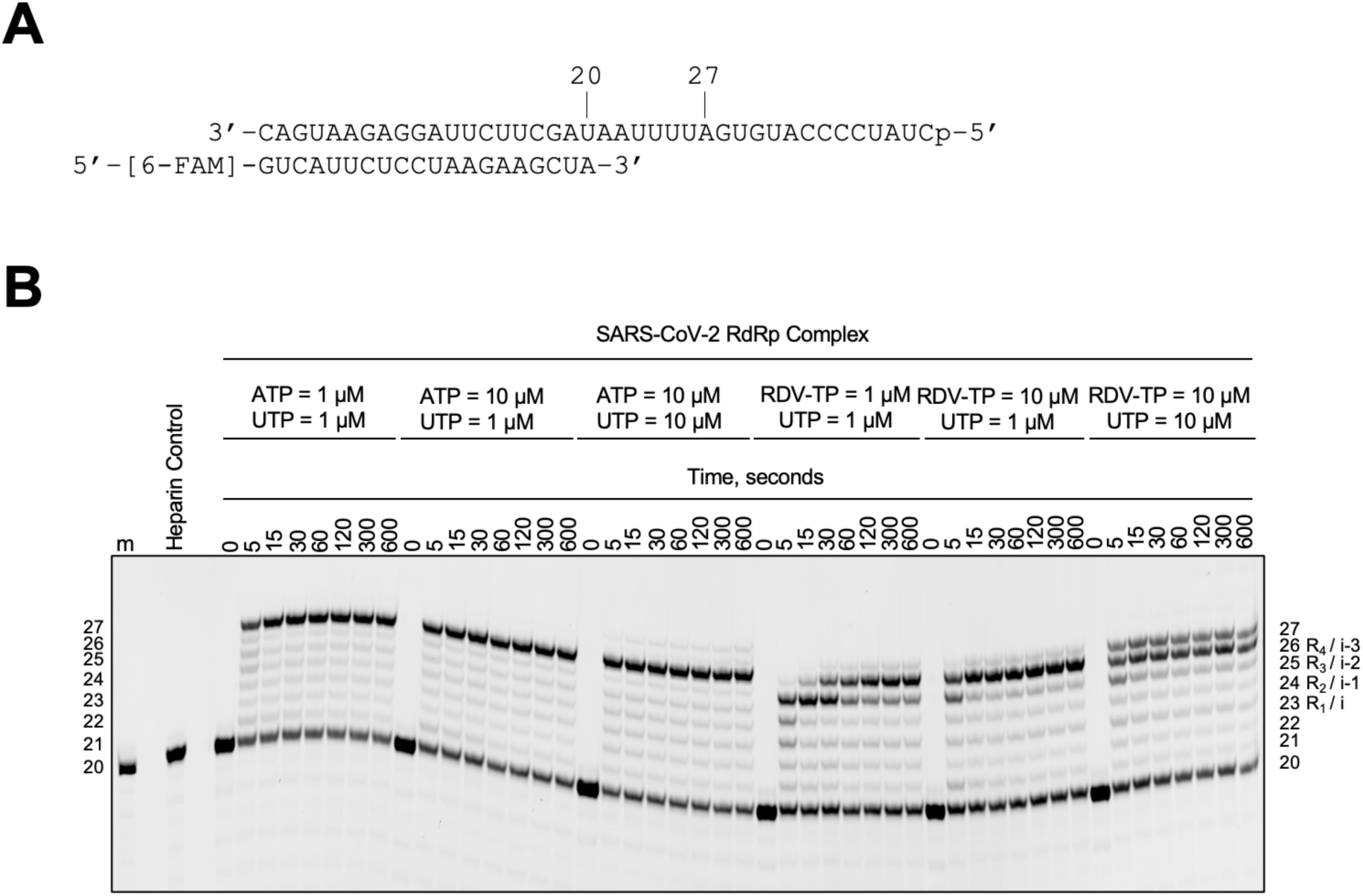
RNA synthesis assay of WT SARS-CoV-2 RdRp complex. **A**, RNA template/primer supporting four successive incorporations of ATP or RTP from positions 23 (i) to 26 (i-3). **B**, Migration pattern of RNA products synthesized by the SARS-CoV-2 RdRp complex. The 5′-6-FAM labelled primer serves as the marker. The *Heparin control* lane is the resulting product when the SARS-CoV-2 RdRp complex is incubated with 4 mg/mL heparin prior to the addition of the RNA duplex and nucleotides. This reaction was equivalent to the longest time point tested (600 seconds). No product formation indicates the concentration of heparin used is sufficient to monitor RNA synthesis under single-turnover conditions.

### Structural impact of the S759A mutation on NTP incorporation and RNA translocation

S759 is in the conserved catalytic hairpin S759-D760-D761 (SDD) of motif C and is unique to the *nidovirales* order; other positive-sense RNA virus RdRps have a glycine (GDD) in the place of serine in the active-site hairpin^32^. Therefore, this active site residue has also been implicated in the processivity and accuracy of RNA synthesis by the coronavirus RdRp that is necessary for the replication of its genome^33,34^. A previous mutagenesis study found that the S759G substitution in SARS-CoV-2 RdRp reduced the fidelity of RNA synthesis^33^; the authors hypothesized that the improved fidelity relates to the loss of an essential interaction between S759 and the 2′-OH of the nucleotide at the 3′-end of the primer.

We compare the structures of S759A-RMP and S759A-AMP complexes with their respective wild-type complexes. The primer 3′-end in the S759A-RMP complex is locked at the N-site following three RTP incorporations, i.e., the structure comparison also did not show any distinct impact of the mutation on the binding mode or conformation of the three incorporated RMPs. The S759A mutation is expected to impact the positioning of the primer 3′-end at the P-site. In fact, our S759A-AMP complex with its primer 3′-end nucleotide ribose ring positioned over the mutation site reveals an important difference from the WT-AMP structure. In the WT-AMP structure, the ribose ring of the primer 3′-end nucleotide has a 3′-endo conformation, and the 2′-OH group is recognized by the H-bond interaction with the Oγ atom of S759, while the ribose ring conformation is stabilized as 3′-endo by this interaction (Figure 5A). This interaction with the 2′-OH group with S759 Oγ is important for the fidelity of the RdRp. In contrast, the H-bond is lost in the S759A-AMP structure. Interestingly, in the mutant complex, the ribose ring at the primer 3′-end adopts a 2′-endo conformation and is also shifted by approximately 1 Å compared to the wild-type complex (Figure 5B). Consequently, the primer terminal 3′-OH in the mutant structure is shifted by ∼1.3 Å, and 2′-OH is shifted by ∼2.3 Å compared to their respective positions in the WT-AMP structure. Notably, the LADD/LSDD active-site hairpin conformation remains unaffected by the S759A mutation, regardless of the observed shift in the primer 3′-end i.e, the mutation repositions the primer 3′-end nucleotide with respect to the polymerase active-site and the structural differences between the WT-AMP and S759A-AMP provide an explanation for the observed lower fidelity of the S759G mutant RdRp described above.

**Figure 5:**
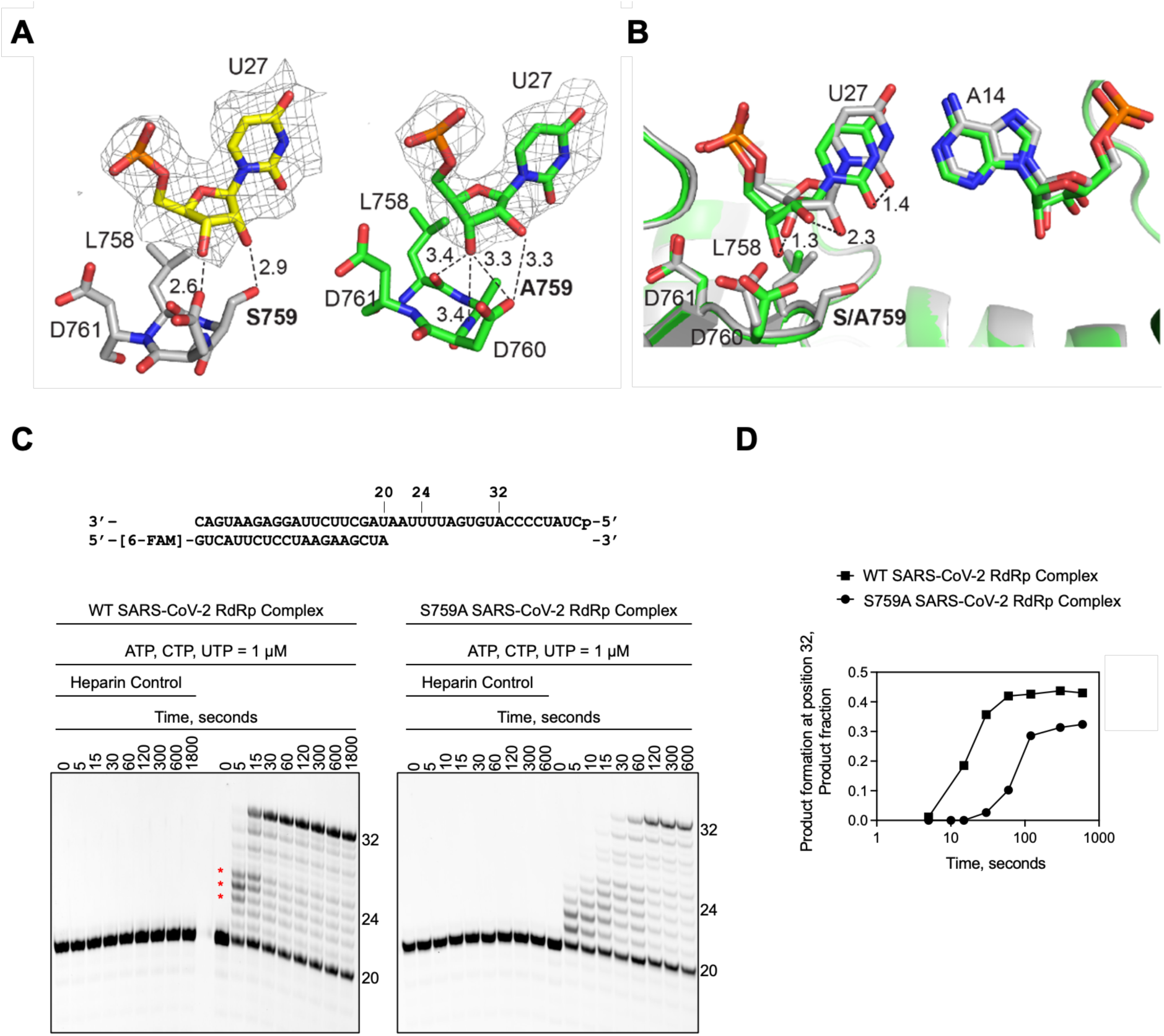
Impact of the S759A mutation on the substrate interaction, and processive RNA synthesis under single turnover conditions by WT and S759A SARS-CoV-2 RdRp complex. **A**, Hydrogen bond network of the primer 3′-end nucleotide at P-site in the WT-AMP (left, grey) and in S759A-/AMP (right, green) structures. The active site hairpin residues (LSDD/LADD) and the U27 are depicted as sticks, with all the distances below 4.0 Å indicated as dashed lines. **B**, Overlap of the P strand 3′-OH between WT-AMP (grey) and S759A-AMP (green) structures. The U:A bp and the LSDD/LADD residues are shown as sticks. The shift of U27 between the two structures is indicated as dashed lines with the distance annotated in Å. **C**, The RNA template/primer supporting RNA synthesis in our experiment is at the top. The addition of ATP, CTP, and UTP enables the synthesis of a 32-mer product. Below is the migration pattern of RNA products synthesized by the SARS-CoV-2 RdRp WT (*left*) or S759A (*right*) complex. The *Heparin control* block is the resulting product when the SARS-CoV- 2 RdRp complex is incubated with 4 mg/mL heparin prior to the addition of the RNA duplex and nucleotides. No product formation at any time point indicates the concentration of heparin used is sufficient to monitor RNA synthesis under single-turnover conditions. **D**, Quantification of the RNA synthesis reaction products in **C** shows the lower catalytic efficiency of the mutant RdRp.

Interestingly, we observed a structural aberration in loop 617–624 of the palm domain, a part of the N-site (Figure S5). This structural difference is observed in both S759A-AMP and S759A- RMP complexes. i.e., the structural change in this mutant is observed irrespective of whether the N-site is occupied or empty, and may have implications for associated mutations, such as V792I^35^. Several RDV-resistant mutations in Nsp12 have been identified both clinically and *in vitro*. The conferred resistance by different Nsp12 mutations is, P323L/V792I (2.2-fold), C799F (2.5-fold), K59N/V792I (3.4-fold) and E802D (∼6-fold)^36–40^. Two independent *in vitro* studies identified that the S759A substitution in Nsp12 confers ∼10-fold resistance to RDV^34,35^. Moreover, the analogous mutation in murine hepatitis virus, an orthologous coronavirus, shared this resistant phenotype^35^.

### S759A mutant is catalytically less efficient than the wild-type RdRp

To address the impact of the mutation, we evaluated processive elongation by the WT and S759A complex (Figures 5C and D). These reactions were performed under single-turnover conditions, as discussed earlier, and initiated by the addition of ATP, CTP, and UTP. For the WT RdRp complex, intermediate products of ∼24-nt, form at earlier reaction timepoints (5 and 15 seconds) before they are extended to the anticipated 32-nt RNA product (Fig. 5C, *left, red stars*). Due to the presence of heparin, we can be certain that this observation is not due to the dissociation of the enzyme but instead suggests that the WT RdRp complex changes speed during RNA synthesis. Notably, the 24-nt intermediate product would correspond to the i-4 position detailed in the RNA duplex backbone evaluation (Figure 3). Therefore, this change in speed could be in response to the natural RNA flexibility observed at position i-4. For the S759A RdRp complex, RNA catalysis is slower compared to the WT (Figure 5C, *right*). In this instance, product formation does not appear to plateau until 120 seconds, approximately 90 seconds later than the WT complex (Figure 5D). This observation aligns with a previous study, which found that the S759A complex demonstrated increased stalling during RNA synthesis^34^. Together, these structural and biochemical findings help explain the role of S759 in both fidelity and elongation. Moreover, we provide the evidence that 1) the RNA flexibility impacts the speed of RdRp RNA synthesis, and 2) a reduced processive elongation by the S759A RdRp complex is likely responsible for the reduced viral fitness observed in cell culture^35^.

## Discussion

### Site-specific preference for RTP incorporation

To elicit an antiviral effect, NAs must efficiently compete with natural NTP pools for incorporation by a viral polymerase. However, several factors can diminish an NA’s efficacy, which are often not captured in simplified biochemical assays. Here, we explored the implications of competitive ATP substrate availability on RTP incorporation. ATP concentration in the cell has been reported to be as high as 3 mM^41^. However, its exact concentration in different cellular compartments remains unknown, making it difficult to assess the true competition between RTP and ATP for the RdRp active site. Nonetheless, ATP concentration is likely much higher than RTP. Even though biochemically, RTP incorporation is preferred over ATP^21^, the comparatively higher concentration of ATP may limit the number of RTP incorporations during minus-strand RNA synthesis. This, in turn, will ultimately reduce the number of RMPs as templating base during replication. The preferential incorporation of first RTP, even at 1/10^th^ of its relative concentration compared to ATP, as observed in our competition assay, reflects the broader efficacy of remdesivir against a wide range of viral RdRps; i.,e, RTP, as a substrate, is superior to ATP under single-incorporation conditions. Structurally, the ribose ring of RTP appears to fit better to the N-site, and the structural rigidity of the ring over the ribose ring of ATP may entropically favor RTP binding over ATP to an empty N-site (Figure 4C). However, for subsequent binding, RTP reduces the incorporation efficiency primarily because the next NTP/RTP requires the translocation of the primer 3′-end nucleotide out of the N-site, a step in which RTP may have a disadvantage because of its less adaptable sugar ring and the C1’ cyano group.

### Potential RMP sites in a synthesized negative-strand RNA

To evaluate the greater impact of the translocation deficiency introduced by the rigid R:U bp, we considered the SARS-CoV-2 genome (accession: NC_045512.2) and the number of opportunities for RTP incorporation during transcription. Out of the approximately 30 kb genome, UMP comprises 9,594 bases; however, 2,439 of these sites are followed by a subsequent UMP which will favor ATP incorporation and, likely reducing the chances for RTP incorporation by approximately 25%. Furthermore, our data show that RTP incorporation is also hindered when RMP is at the i-2 and i-3 positions. There are 5,597 instances where one or two bases separate two UMPs in the viral genome. Overall, the sequence-dependent translocation effect could decrease the opportunities for RTP incorporation by up to 84% when the ATP concentration is 10x or higher than RTP. It is also known that an exonuclease-deficient coronavirus is more susceptible to RDV treatment, indicating that at least some RMP residues are excised during replication^11^. Consequently, the presence of the proofreading exonuclease would likely amplify the effect of reduced opportunities for RTP incorporation. Together, these observations provide a holistic perspective on RTP, encompassing how its chemical modifications, although necessary for its antiviral effect, introduce rigidity that may influence subsequent incorporation events. These lessons should be considered when developing and evaluating future non-obligate NTP analogs targeting RNA viruses.

### The structural implications of S759A on resistance to RTP

The S759A substitution, which confers resistance to RDV, was initially identified through serial passaging of SARS-CoV-2 *in vitro*^30^. To date, this mutation has not been identified in the COVID-19 patient population that has shown resistance to RDV treatment. However, previous biochemical and structural studies have demonstrated that S759 plays an integral role in the efficient incorporation of RTP^24,34,35^. The S759A complex demonstrated a 5- to 10-fold reduction in RTP incorporation. In the mutant structure, we observe that the 3′-end RMP redefines its interactions with surrounding protein residues in the N-site, and the number of interactions is reduced (Figure 6). This reduced interaction is primarily caused by the reorganization of the Nsp12 617–624 loop away from the RMP (Figure S5). The structural reorganization might weaken the stabilization of RMP, which may partially explain the lower affinity of this mutant for remdesivir. The primer 3′-end nucleotide at P-site in WT-ATP structure is significantly impacted by the S759A mutation (Figure 5A), and the repositioned 3′-end appears to be less favorable for next nucleotide addition. This shift in 3′-OH position is likely to reduce the catalytic efficiency of the mutant and further decrease the translocation efficiency of RMP at the primer end, which could increases the chance of excision^34^.

**Figure 6:**
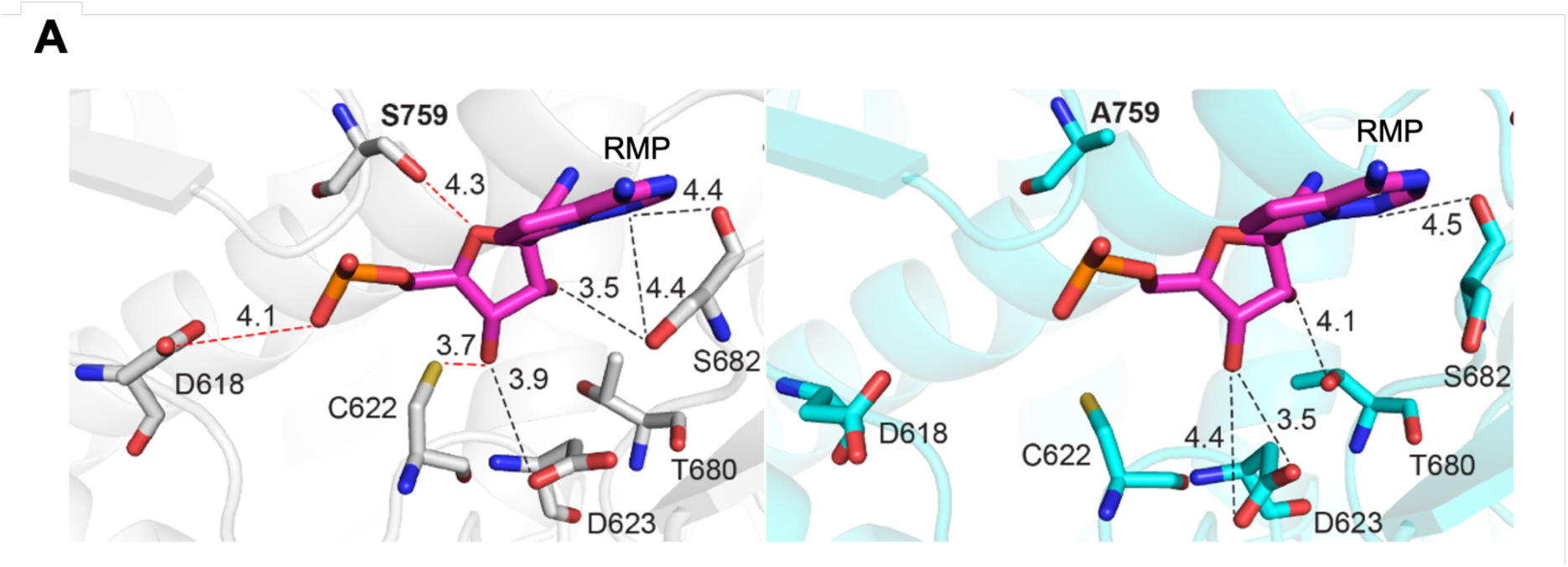
The interactions of RMP with N-site in the wild-type and S759A mutant SARS-CoV-2 RdRp complexes. **A**, Interactions of RMP (magenta) in the N site in the WT-RMP (left, grey) and S759A- RMP (right, cyan) structures. For clarity, only RMP and the interacting residues are shown. Distances are in Å and indicated by dotted lines; the red dotted lines indicate the interactions that are lost in the S759A mutant and in black the ones that differ between the two structures.

### SARS-CoV-2 Nsp12 S759A vs HIV RT M184V/I mutations

In HIV RT, a viral DNA polymerase, M184 is the positional equivalent of S759 in SARS-CoV-2 Nsp12. The RT mutation M184V/I confers resistance to nucleoside drugs containing a L-ribose ring, such as 3TC and FTC^42^. Structures and biochemical studies have explained that the M184V/I mutation introduces a wider bifurcated sidechain that results in a steric conflict with the altered L-ribose ring conformation of 3TC/FTC^43^, and hepatitis B (HBV) DNA polymerase also develops 3TC/FTC resistance by a similar mechanism^44^. Conversely, S759A resistance is caused by a loss of interaction, which is likely in response to the very efficient rates of incorporation for RTP. The contrast between these mechanisms of resistance provides an elegant example of how altering an active site residue can confer resistance through different molecular mechanisms. Moreover, this observation exemplifies that understanding the structural and biochemical properties of a given analog can help understand antiviral resistance mechanisms.

## Materials and methods

### Protein cloning, expression and purification

Plasmid pRSFDuet-1 (nsp8-nsp7)(nsp12) expressing the wild-type form of SARS-CoV-2 RdRp was obtained from Addgene (ref. 165451). The S759A point mutation was introduced by PCR using 2 overlapping primers (P1_S759A 5′- GGCATCATCTGCCAGAATCATCATGCTGAAG-3’ and P2_S759A 5′- ATGATTCTGGCAGATGATGCCGTTGTTTGC-3’). One ng of wild-type plasmid was used as template, and the PCR reaction was performed by adding 0.5 μL of Q5 High-Fidelity DNA Polymerase (New England Biolabs) following the manufacturer’s protocol in a final volume of 50 μL. PCR products (5 μL) were loaded on 1% TAE agarose gel for quality control, then 20 μL were digested with Dpn1 (Thermo Scientific) for 1h at 37 °C. Ten μL of the digested product was used to transform DH5α *E. Coli* (Invitrogen) for plasmid selection and amplification. The presence of the S759A point mutation was confirmed by sequencing (Macrogen). Both wild-type and S759A mutant of SARS-CoV-2 RdRp were expressed and purified as described in Madru et al., 2021^45^.

### RNA synthesis assay

The template and primer used in the gel-based RNA synthesis studies were 5′-phosphorylated and purchased from Dharmacon. The template (5′- CUAUCCCCAUGUGAUUUUAAUAGCUUCUUAGGAGAAUGAC-3’) and primer (5′-6-carboxyfluorescein-GUCAUUCUCCUAAGAAGCUA-3’) were annealed by mixing at a 1:1.5 molar ratio (primer to template) in buffer (50 mM Tris-HCl pH 8 and 50 mM NaCl) and heating to 95℃ before allowing to cool to room temperature.

All reactions were performed at room temperature using the purified SARS-CoV-2 complex described above. Briefly, 2 µM RdRp complex and 100 nM RNA substrate were incubated in reaction buffer (25 mM Tris-HCl pH 8, 5 mM MgCl_2_) for 15 minutes. RNA synthesis was initiated by the addition of nucleotides (various concentrations) and heparin (4 mg/mL) and stopped using equal parts formamide/EDTA (50 mM). The reaction products were incubated at 95°C for 5 minutes and then resolved by 20% Urea-PAGE, and the fluorescent signal was scanned using Typhoon PhosphorImager (Cytiva). The data were analyzed using GraphPad Prism 7.0 (GraphPad Software, Inc., https://www.graphpad.com).

### Mass Spectrometry

Generation of the dsRNA for MS analysis was performed similarly to the RNA synthesis assay described above, with slight variations. Here, 2 µM RdRp complex and 10 µM RNA substrate were incubated in the reaction buffer for 15 minutes. RNA synthesis was initiated by the addition of nucleotides (various concentrations) and allowed to proceed for approximately 16 hours. The reaction was stopped with an equal volume of phenol-chloroform. The aqueous layer was then passed through a Micro Bio-Spin® P-30 column (Bio-Rad, #7326223) that was equilibrated with H_2_O. The collected flow-through underwent precipitation with sodium acetate (0.3 M final) and isopropanol before being washed twice with 70% ethanol. The pellet was resuspended in 10 µL Milli-Q H_2_O to achieve a final concentration of ∼15 µM. The RNA synthesis products were then subjected to MALDI-TOF MS with a 2,4,6- trihydroxyacetophenone (THAP) matrix (Dr. Randy Whittal and Dr. Joseph Utomo, Mass Spectrometry Facility, Edmonton, AB, Canada).

### Assembly of RdRp/RNA complexes and substrate incorporation for cryo-EM

Template (5′- CUAUCCCCAUGUGAUUUUAAUAGCUUCUUAGGAGAAUGAC-3’) and primer (5′- GUCAUUCUCCUAAGAAGCUA-3’) RNAs were purchased from Integrate DNA technologies, resuspended separately in H_2_O and mixed in a 1:1 molar ratio in TE buffer. The mixture was incubated at 60 °C for 5 minutes then cooled down at room temperature to favor prime-template annealing. RdRp (wild-type or S759A) purified by anion exchange (HiTrap Q HP, Cytvia) was incubated with annealed RNA (in a complex:RNA molar ratio of 1:1.5) and UTP (50x molar excess to RNA) to incorporate the first two nucleotides on the primer strand, in a buffer containing 20 mM HEPES pH 7.5, 75 mM NaCl, 5 mM MgCl_2_ and 5 mM DTT. Complex was incubated at room temperature for 10-15 min and then purified by SEC on a Superdex 200 Increase 10/300 GL column connected to an AKTA PURE 25 FPLC system (GE Healthcare Life Sciences) maintained at 6 °C. Fractions of interest were pooled and concentrated using a 10-kDa-cutoff Amicon ultra-centrifugal filter (Merck Millipore) and stored at −80 °C prior further use. The RdRp/RNA complex (wild-type or S759A) purified by SEC was used for substrate incorporation by adding either ATP or Remdesivir triphosphate (100x molar excess to RNA) and UTP (50x molar excess to RNA). Samples were incubated at room temperature for 10-15 minutes then kept on ice prior grid preparation.

### Cryo-EM grid preparation and data collection

Vitrification of RdRp/RNA complexes was done on Quantifoil R 1.2/1.3 holey carbon grids (Au300) using a Leica EM GP (Leica Microsystems). The grids were glow-discharged for 45 s at 25 mA current with the chamber pressure set at 0.3 mbar (PELCO easi-Glow; Ted Pella). Glow-discharged grids were mounted in the sample chamber of a Leica EM GP at 8 °C and 95% relative humidity. Optimized grids for all samples were obtained by pipetting 3 μL of sample at ∼3 μM. The sample (in 20 mM HEPES pH 7.5, 75 mM NaCl, 5 mM MgCl_2_, 5 mM DTT) was incubated on the grids for 10 s before back-blotting for 12 s using two pieces of Whatman Grade 1 filter paper, and the plunge-freezing was carried out by dipping the grids in liquid ethane at a temperature of −172 °C. The grids were clipped and mounted on a 200-keV Glacios cryo-transmission electron microscope (Thermo Fisher) with an autoloader and Falcon 4i direct electron detector equipped with a Selectris energy filter as installed in our laboratory. High-resolution datasets were collected on the Glacios using EPU software version 3.5.1 (Thermo Fisher). Electron microscopy data were recorded as movies in counting mode at a nominal magnification of ×130.000 yielding a pixel size of 0.9 Å. The total exposure time was 8.13 s for WT-AMP, 8.18 s for WT-RMP and S759A-AMP and 15.56 s for S759A-RMP for a total dose of 40 e^-1^/A^2^ for all datasets, fractionated to 2043 frames for WT-ATP, 2052 frames for WT-RMP, S759A-AMP and S759A-RMP. The data collection parameters for all structures are listed in Supplementary Table 1.

### Cryo-EM data processing

An overview of the workflow used for processing each structure is given in Figure S2. For each dataset, individual video frames were motion-corrected and aligned using MotionCor2^46^ as implemented in the Relion 4 package^47^ and the contrast transfer function parameters were estimated by CTFFIND-4^48^. The particles were automatically picked using the reference-free Laplacian-of-Gaussian routine in Relion 4. The picked particles were cleaned by cycles of two-dimensional (2D) and 3D classifications. A density map for RdRp (EMD-30210) was blurred to 40 Å resolution and used as the reference for initial 3D classifications to eliminate partially disordered and incomplete particles. The homogeneous particle sets from the best 3D class were used to calculate gold-standard 3D auto-refined maps and corresponding masks. For each structure, the particles were repicked at the 3D auto-refinement stage and classified further within its mask. Particles were extracted with a box size of 256 pixels. Final 3D classification generated a distinct class (Figure S2). The final set of particles for each structure was used to calculate gold-standard auto-refined maps, further improved by Bayesian polishing and contrast transfer function refinement. All data processing steps were carried out using Relion 4. The local-resolution maps were calculated using ResMap and the orientation plots were generated by Relion 4.

### Model building

Data processing yielded 2.8 Å, 2.9 Å, 3.1 Å, and 2.9 Å density maps for WT-AMP, WT-RMP, S759A-AMP and S759A-RMP, respectively, and these were used to fit the atomic models for the respective structures. Previously published SARS-CoV-2 RdRp in complex with RNA (PDB ID: 7DOK) was used as reference for building the model of the WT-AMP structure and then this model was used as initial reference for the other three structures. All model building was carried out manually using COOT^49^ coupled with iterative rounds of real-space structure refinement using Phenix 1.19.2^50^. In all structures a single copy of Nsp8 could be modelled with residues 191-198 of this subunit missing from all four structures. In addition, residues 1-75 are missing from WT-AMP, 1-77 from WT-RMP, 1-76 from S759A-AMP and 1-77 from S759A-RMP structures. Nsp7 residues 63-83 are missing in WT-AMP and S759A-AMP, 1 and 62-83 in WT-RMP, 1 and 66-83 in S759A-RMP structures. Nsp12 missing residues are the following: 1-5, 107-110, 898-912 and 931-932 in the WT-AMP structure; 1-28, 53-66, 99-114, 898-912 and 931-932 in the WT-RMP structure; 1-6, 107-110, 898-912 and 931-932 in the S759A-AMP structure; 1-28, 55-68, 100-114, 898-912 and 931-932 in the S759A-RDV structure. All structure figures were generated using PyMol (https://pymol.org/2/), Chimera^51^ and ChimeraX^52^.

## Acknowledgements

The authors acknowledge Dr. Abhimanyu Singh for advice on protein expression, Dr. Randy Whittal and Dr. Joseph Utomo at the Mass Spectrometry facility at the University of Alberta for the mass spectrometry analysis.

## Author Contributions

Conceptualization: K.D., C.J.G., M.G.; Investigation and Methodology: C.J.G. and H.W.L. Biochemistry, C.J.G. Mass Spec; M.F.O. protein purification; M.F.O., Q.G., and B.D-W. cryo-EM and structural studies; Data analysis: C.J.G., H.W.L., M.G. biochemistry, C.J.G., K.D. Mass spec; M.F.O., Q.G., and K.D. Structural biology; Funding acquisition: K.D. and M.G.; Project administration and supervision: K.D. Original draft written by: C.J.G., M.F.O., K.D., & reviewed and edited by all authors.

## Data Availability

The data that support this study are available from the corresponding author upon reasonable request. The coordinates and cryo-EM density maps for the structures WT-AMP, WT-RMP, S759A-AMP, and S759A-RMP are deposited with PDB accession codes/EMDB codes 9SAR/EMD-54695, 9SAP/EMD-54693, 9SAQ/EMD-54694, and 9SAO/EMD-54692, respectively.

## Supplementary Data

Supplementary data are available online, containing Table S1 and Figures S1 – S5.

## Supplementary Information

**Table S1.** Single particle cryo-EM data and structure analysis statistics.

|  | WT-AMP<br>(EMD-54695)<br>(PDB 9SAR) | WT-RMP<br>(EMD-54693)<br>(PDB 9SAP) | S759A-AMP<br>(EMD-54694)<br>(PDB 9SAQ) | S759A-RMP<br>(EMD-54692)<br>(PDB 9SAO) |
| --- | --- | --- | --- | --- |
| **Data collection and processing** |  |  |  |  |
| Magnification | 130,000x | 130,000x | 130,000x | 130,000x |
| Voltage (kV) | 200 | 200 | 200 | 200 |
| Electron exposure (e-/Å<sup>2</sup>) | 40 | 40 | 40 | 40 |
| Defocus range (um) | -0.6 to -1.8 | -0.6 to -1.8 | -0.6 to -1.8 | -0.6 to -1.8 |
| Pixel size | 0.9 | 0.9 | 0.9 | 0.9 |
| Symmetry imposed | C1 | C1 | C1 | C1 |
| Initial particle images (no.) | 654,460 | 1,237,567 | 1,002,711 | 738,355 |
| Final particle images (no.) | 109,207 | 145,169 | 93,716 | 167,051 |
| Map resolution (Å) | 2.8 | 2.9 | 3.1 | 2.9 |
| FSC threshold | 0.143 | 0.143 | 0.143 | 0.143 |
| Local resolution estimation (Å) | 6.2 to 2.8 | 6.3 to 2.9 | 6.2 to 3.1 | 6.5 to 2.9 |
| **Refinement** |  |  |  |  |
| initial model used (PDB ID) | 7DOK | 9SAR | 9SAR | 9SAR |
| Model resolution (Å) | 3.0 | 2.6 | 2.7 | 2.8 |
| FSC threshold | 0.143 | 0.143 | 0.143 | 0.143 |
| Model resolution range (Å) | 100 to 3.0 | 100 to 2.6 | 100 to 2.7 | 100 to 2.8 |
| Map sharpening B factor (Å<sup>2</sup>) | -94 | -98 | -82 | -105 |
| Model composition |  |  |  |  |
| Non-hydrogen atoms | 9330 | 8663 | 9305 | 8697 |
| Protein residues | 1084 | 1007 | 1082 | 1011 |
| Nucleotide residues | 31 | 25 | 31 | 25 |
| Ligands | Zn:2 | F86:3<br>Zn:2 | Zn:2 | F86:3<br>Zn:2 |
| B factors (Å<sup>2</sup>) |  |  |  |  |
| Protein | 74.86 | 45.04 | 25.78 | 54.58 |
| Nucleotide | 109.08 | 43.81 | 75.59 | 54.18 |
| Ligands | 82.91 | 19.27 | 26.64 | 17.24 |
| R.m.s. deviations |  |  |  |  |
| Bond lengths (Å) | 0.007 | 0.005 | 0.006 | 0.004 |
| Bond angles (°) | 0.937 | 0.793 | 0.873 | 0.618 |
| Validation |  |  |  |  |
| MolProbity score | 1.42 | 1.78 | 1.75 | 1.80 |
| Clash score | 4.30 | 5.46 | 3.21 | 5.97 |
| Poor rotamers (%) | 0.42 | 2.13 | 3.67 | 2.24 |
| Ramachandran plot |  |  |  |  |
| Favored (%) | 96.65 | 96.48 | 96.74 | 96.70 |
| Allowed (%) | 3.17 | 3.52 | 3.26 | 3.30 |
| Disfavored (%) | 0.19 | 0.00 | 0.00 | 0.00 |

**Figure S1:**
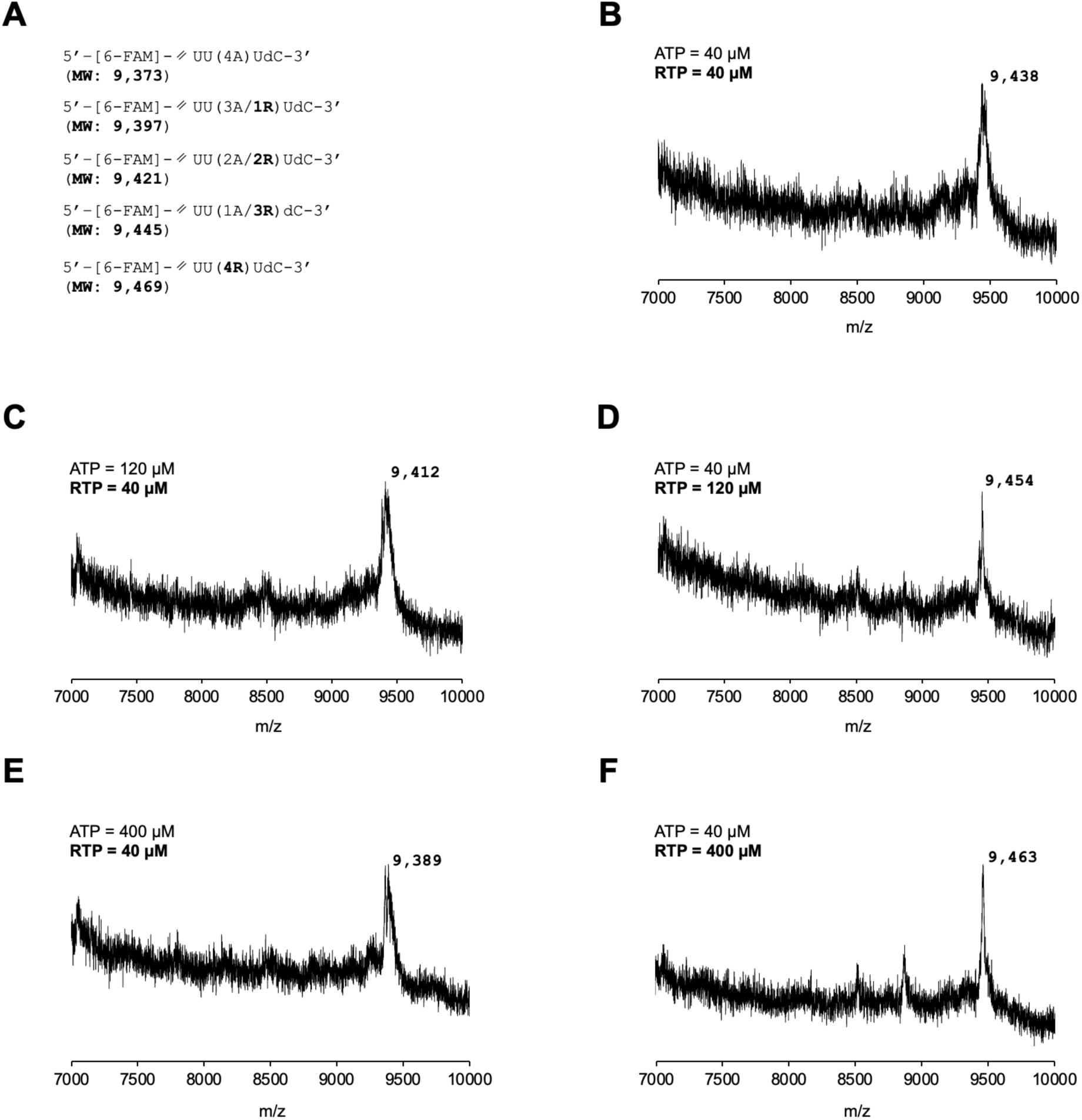
MALDI-TOF mass spectra for the RNA product generated by SARS-CoV-2 RdRp complex under different ratios of ATP to RTP. **A**, the potential sequence outcomes and their predicted molecular mass are shown in brackets below. **B-F** are the RNA MALDI-TOF mass spectra that resulted from different concentrations of ATP and RTP, which are shown above each graph.

**Figure S2A:**
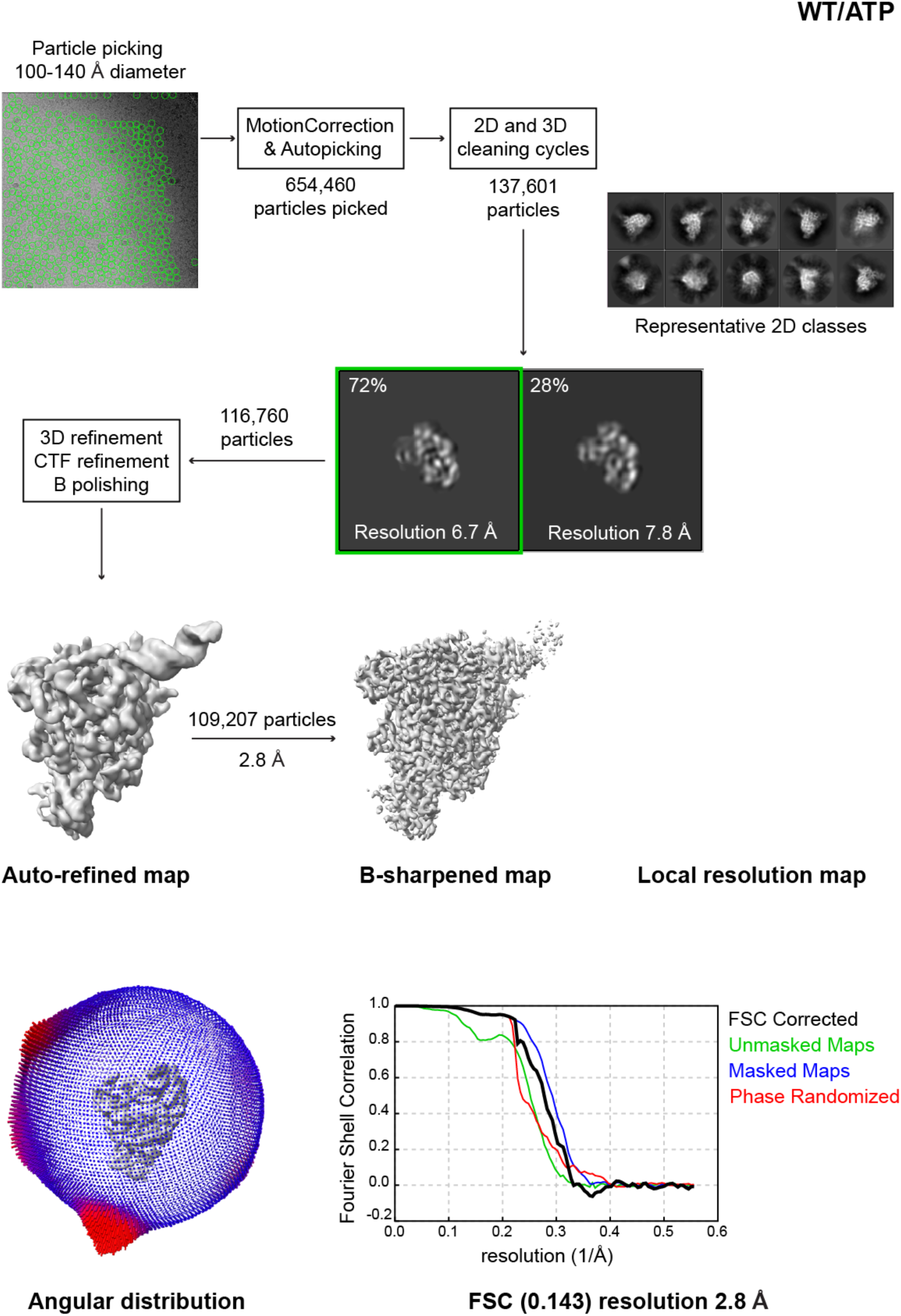
Cryo-EM image processing flowcharts summarizing the data processing and the density map quality, particle distribution and resolution of WT-AMP structure.

**Figure S2B:**
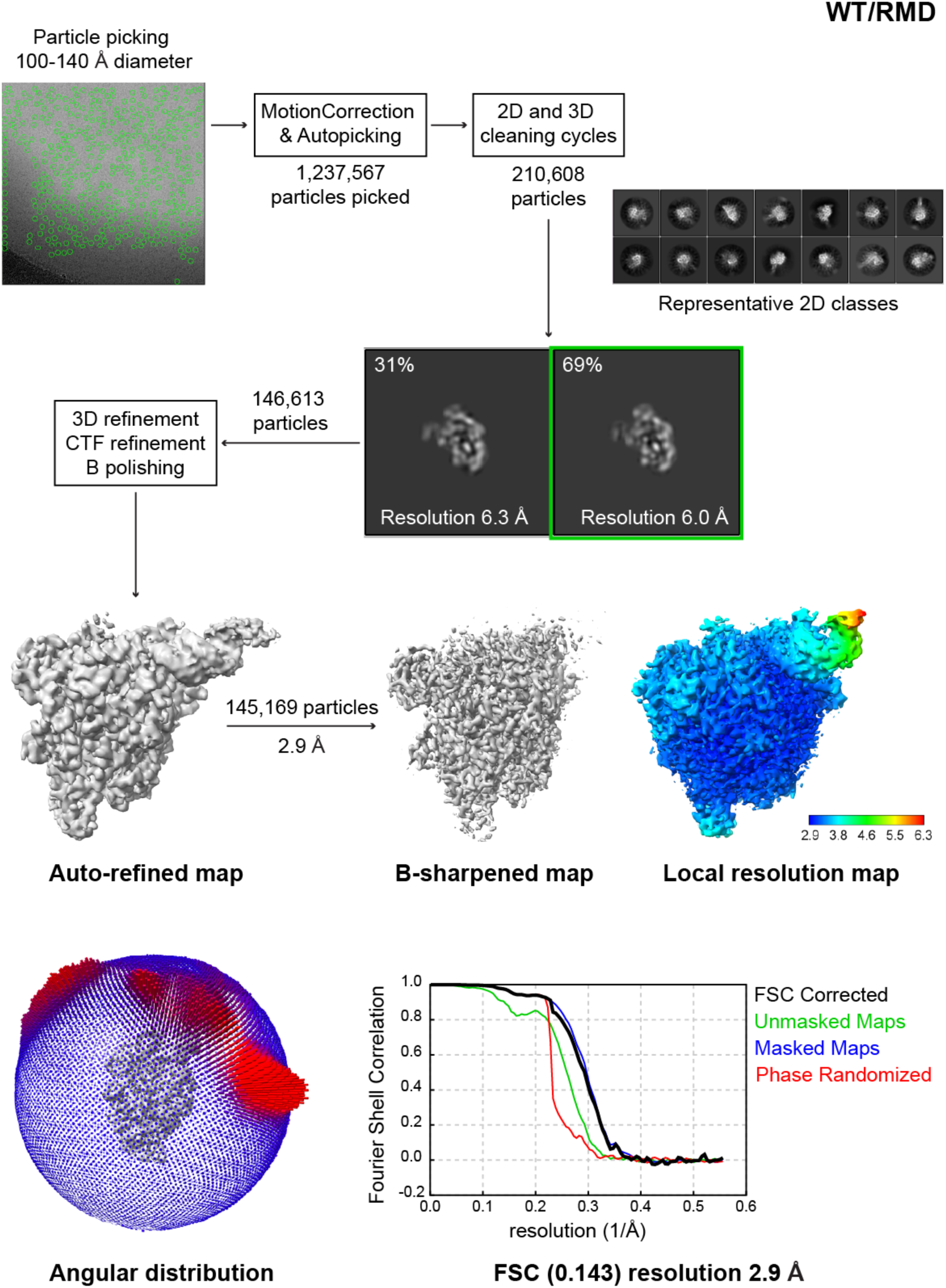
Cryo-EM image processing flowcharts summarizing the data processing and the density map quality, particle distribution and resolution of WT-RMP structure.

**Figure S2C:**
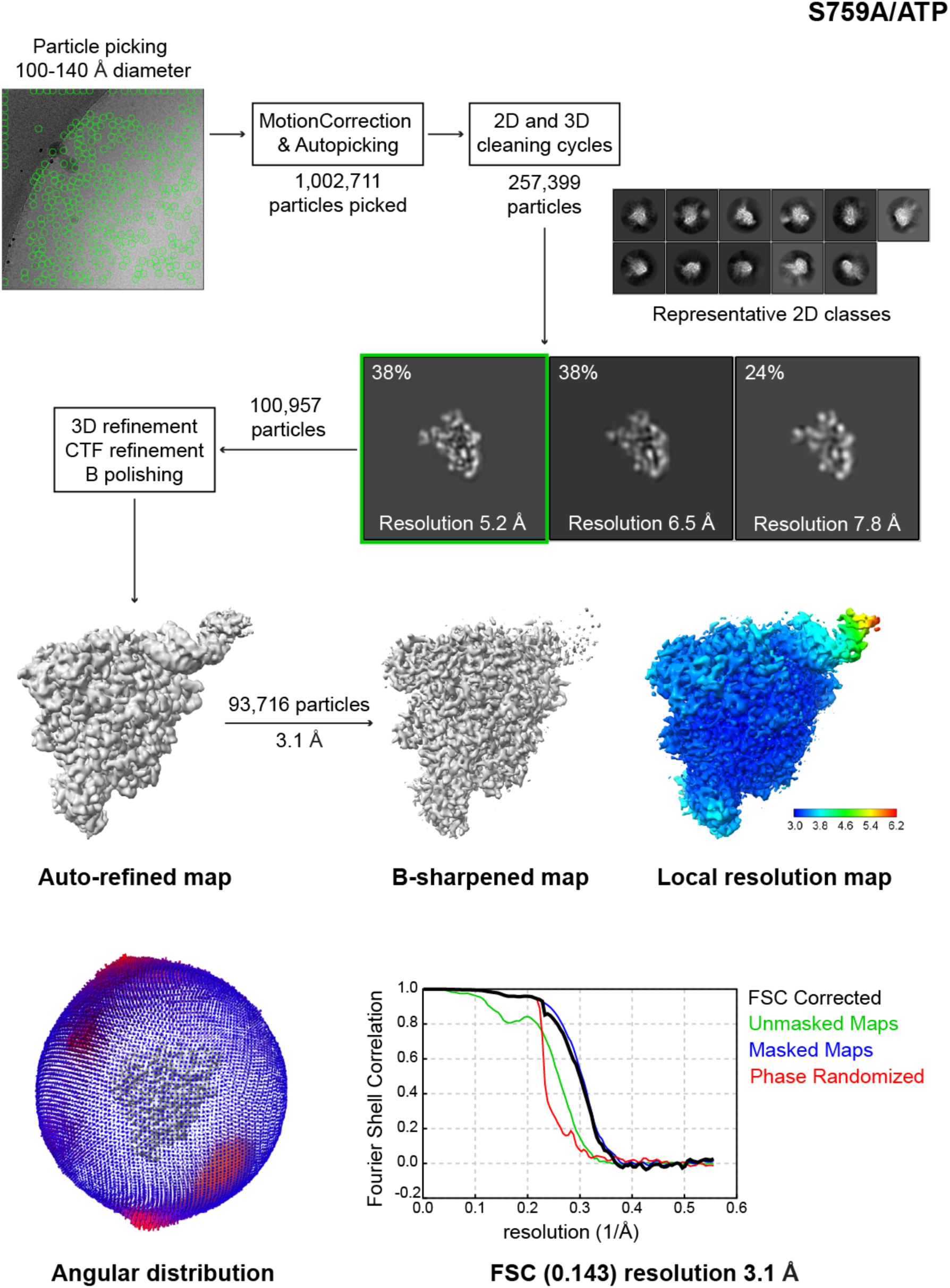
Cryo-EM image processing flowcharts summarizing the data processing and the density map quality, particle distribution and resolution of S759A-AMP structure.

**Figure S2D:**
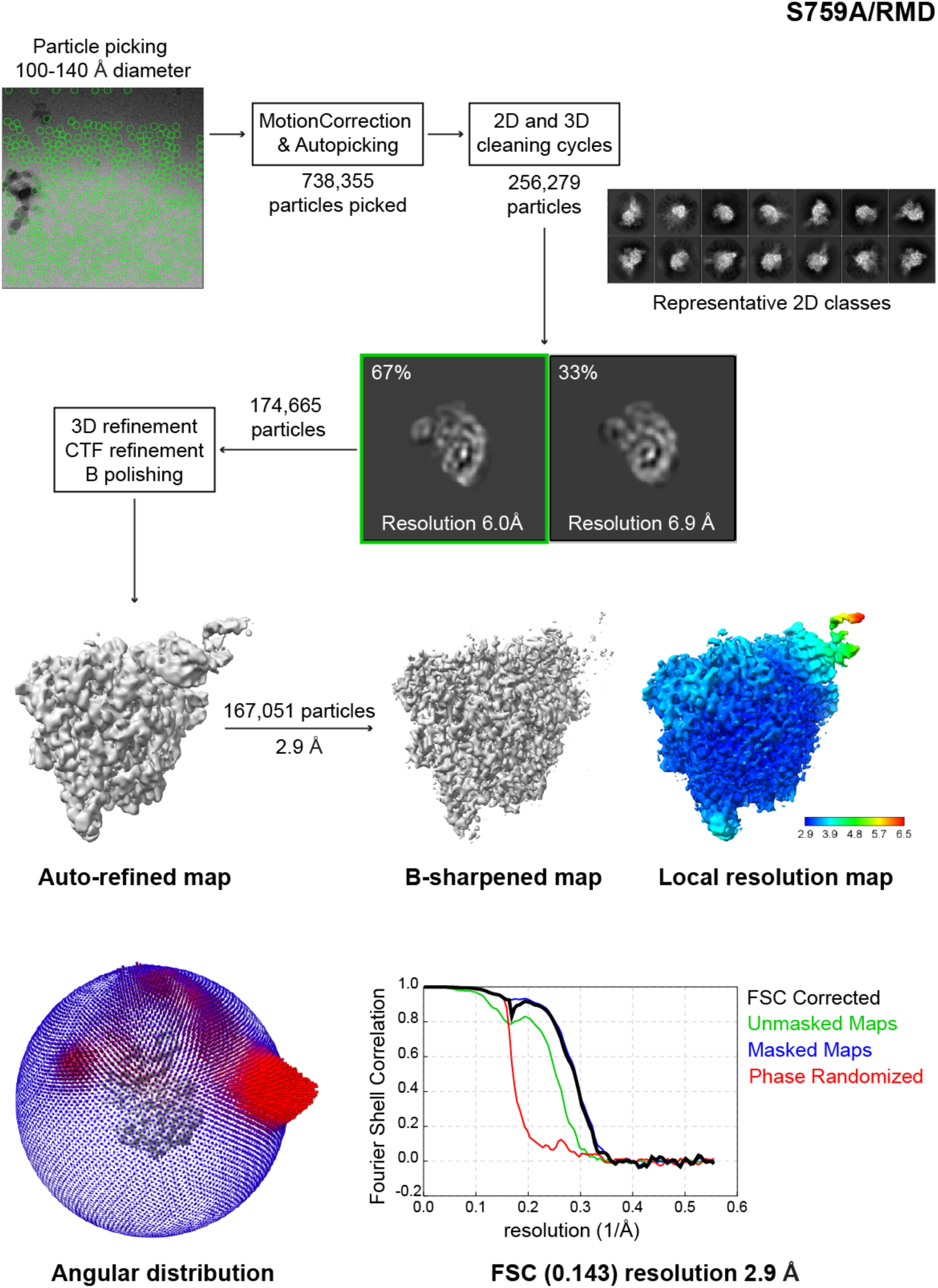
Cryo-EM image processing flowcharts summarizing the data processing and the density map quality, particle distribution and resolution of S759A-RMP structure.

**Figure S3:**
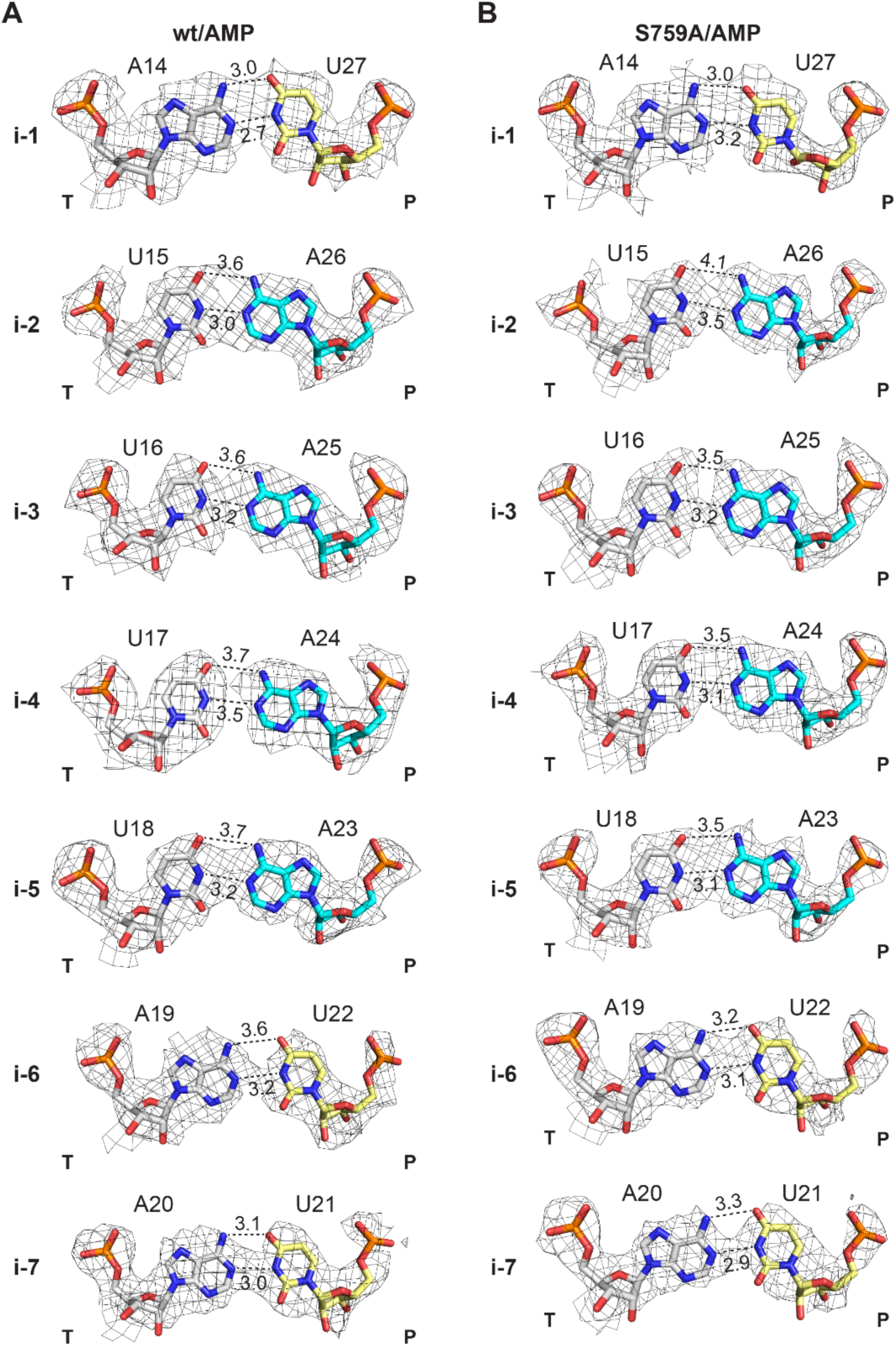
ATP incorporation in WT and S759A RdRp complexes. Atomic model and electron density of different positions of the RNA template(T):primer(P) strands. Bases are shown as sticks, colored following the scheme of Figure 1 and density maps are contoured at 3.0 σ. **A**, WT-AMP complex bps at positions i-7 to i-1 (from bottom to top). **B**, S759A-AMP complex bps modelled at positions i-1 to i-7 (from top to bottom).

**Figure S4:**
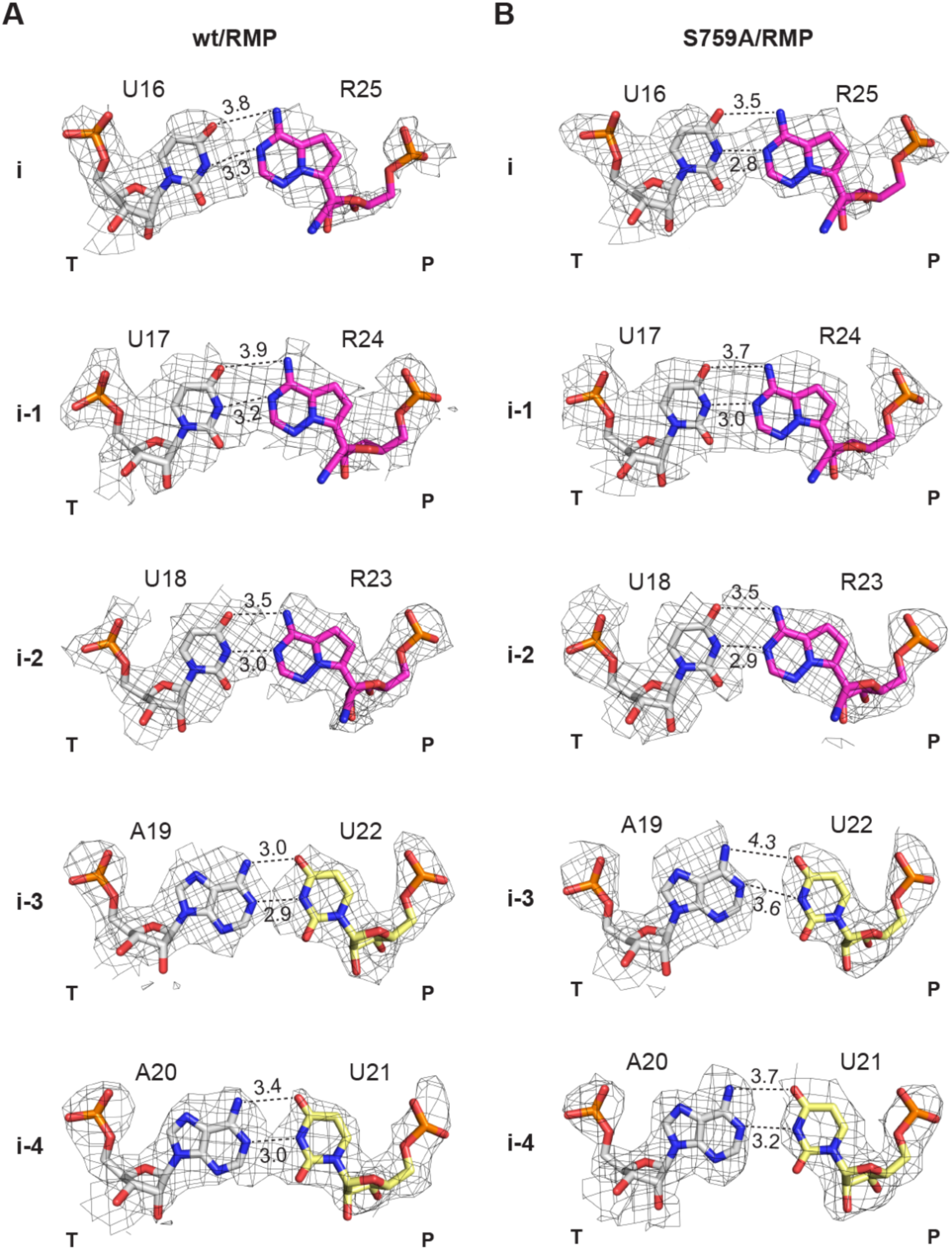
RDV incorporation in WT and S759A RdRp complexes. Atomic model and electron density of different positions in the RNA template(T):primer(P) strands. Bases are shown as sticks, colored as in Figure 1, density maps are contoured at 3.0 σ. **A** WT-RMP complex bpss modelled at positions i-4 to i (from bottom to top). **B** S759A-RMP complex bps modelled at positions i-4 to i (from bottom to top).

**Figure S5:**
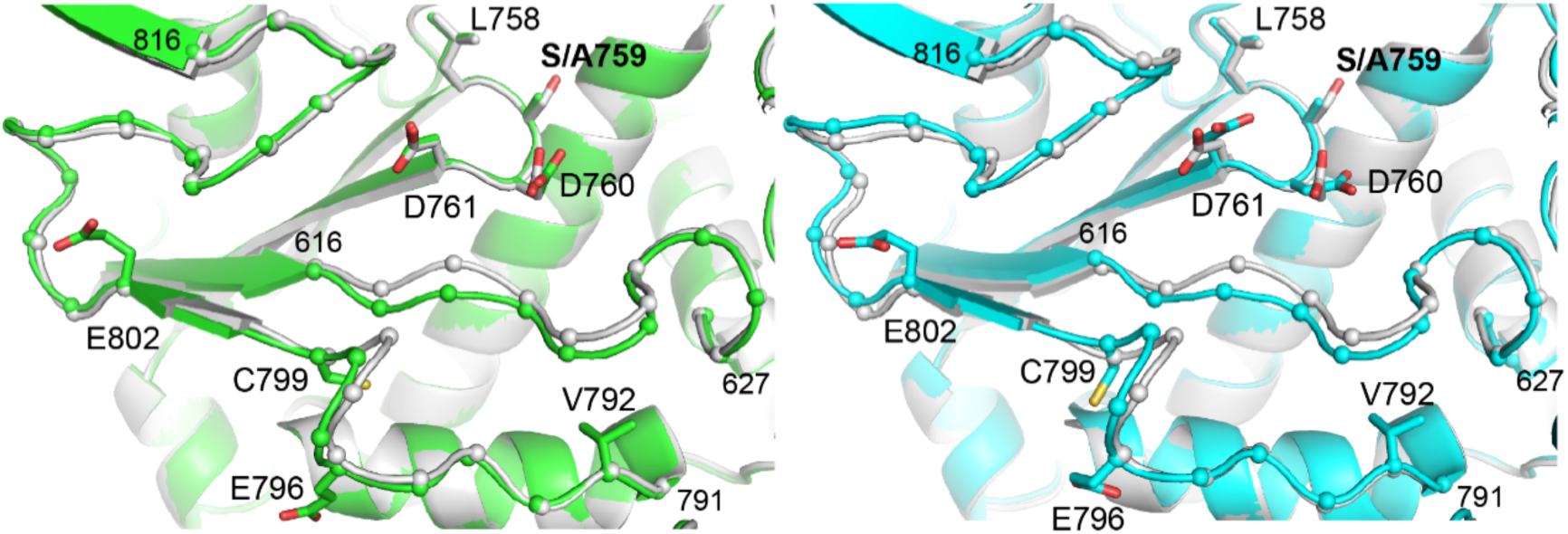
Shift of the palm domain loops in the S759A structures. Overlap of the palm domain segments between the WT-AMP (gray), S759A-RMP (cyan) and S759A-RMP (green). Residues of the LS(A)DD domain are labelled and shown as sticks. The loops showing largest movements are depicted with the C⍺ represented as spheres with only the residues at the extremities numbered for simplicity.

